# Cell-TIMP: Cellular Trajectory Inference based on Morphological Parameter

**DOI:** 10.1101/2024.04.18.590109

**Authors:** Piyush Raj, Himanshu Gupta, Pooja Anantha, Ishan Barman

## Abstract

Cellular morphology, shaped by various genetic and environmental influences, is pivotal to studying experimental cell biology, necessitating precise measurement and analysis techniques. Traditional approaches, which rely on geometric metrics derived from stained images, encounter obstacles stemming from both the imaging and analytical domains. Staining processes can disrupt the cell’s natural state and diminish accuracy due to photobleaching, while conventional analysis techniques, which categorize cells based on shape to discern pathophysiological conditions, often fail to capture the continuous and asynchronous nature of biological processes such as cell differentiation, immune responses, and cancer progression. In this work, we propose the use of quantitative phase imaging for morphological assessment due to its label-free nature. For analysis, we repurposed the genomic analysis toolbox to perform trajectory inference analysis purely based on morphology information. We applied the developed framework to study the progression of leukemia and breast cancer metastasis. Our approach revealed a clear pattern of morphological evolution tied to the diseases’ advancement, highlighting the efficacy of our method in identifying functionally significant shape changes where conventional techniques falter. This advancement offers a fresh perspective on analyzing cellular morphology and holds significant potential for the broader research community, enabling a deeper understanding of complex biological dynamics.

## Introduction

Cell shape is intricately tied to cell function. Previous studies [1] [2] [3] [4] have established that cell shape is controlled by both the physical properties of the plasma membrane and the biochemical reactions involving membrane components and the underlying cytoskeleton. At its core, the shape of cells embodies the collective imprints of intricately interconnected molecular networks, encompassing metabolic, proteomic, epigenomic, and genomic elements [5] [6] [7] [8]. Therefore, it’s unsurprising that clinicians routinely employ cellular morphology for diagnosis, prognosis and treatment of human diseases [9] [10].

Cell morphology has traditionally been quantified on the basis of few geometric parameters on a bright field or fluorescently labelled image [11] [12] [13]. These geometric parameters can broadly be divided into two categories – dimensional (e.g., area, perimeter) and dimensionless (e.g., aspect ratio, shape factor, eccentricity) parameters. The broad morphological parameters have helped unveil several insights and establish links between size, shape, and functionality [14]. Although fluorescent images offer high contrast between the cell and its surroundings and allow organelle-level specificity through dye labeling, it’s important to note that the stained molecules introduced externally can alter the cell’s natural state. Additionally, issues such as phototoxicity and photobleaching can introduce errors in morphological estimations [15] [16]. Therefore, there is a strong desire for contrast agent-free microscopy techniques that enable the study of cells in their native conditions.

Quantitative phase imaging (QPI) is emerging as a prominent tool for extracting quantitative morphological information without the need for external labeling agents [17] [18]. QPI measures the phase shift of light waves as they pass through objects, resulting in the generation of phase images that offer superior contrast compared to other label-free imaging methods. The dimensional and dimensionless parameters from QPI images have helped in label-free cell population monitoring, providing crucial insights into the dynamics of red blood cells [19], cell-nanoparticle interactions [20] and cellular phenotypes during adipocyte differentiation [21]. The wealth of structural information provided by QPI had helped in providing classification ability between cell cycle stage [22], differentiation between cancerous and non-cancerous cells [23] and label-free SARS-CoV-2 detection and classification [24].

Traditionally, most of the cellular morphological analysis methods correspond to classification algorithms or binning into shape modes to categorize into their pathophysical states [13] [25] [26] [23] [27]. Nonetheless, various biological processes, such as cell differentiation, immune responses, and the progression of cancer, undergo a continual and asynchronous shift between cellular states [28] [29] [30]. In this context, traditional analytical techniques often lack the efficacy to elucidate the dynamic essence of these intricate phenomena. This challenge is metaphorically depicted by Waddington’s epigenetic landscape (Fig 1c) [31]. While helpful in analysis, similar values in geometric parameters can still result in large heterogeneity in cell shape [32].

**Figure 1:**
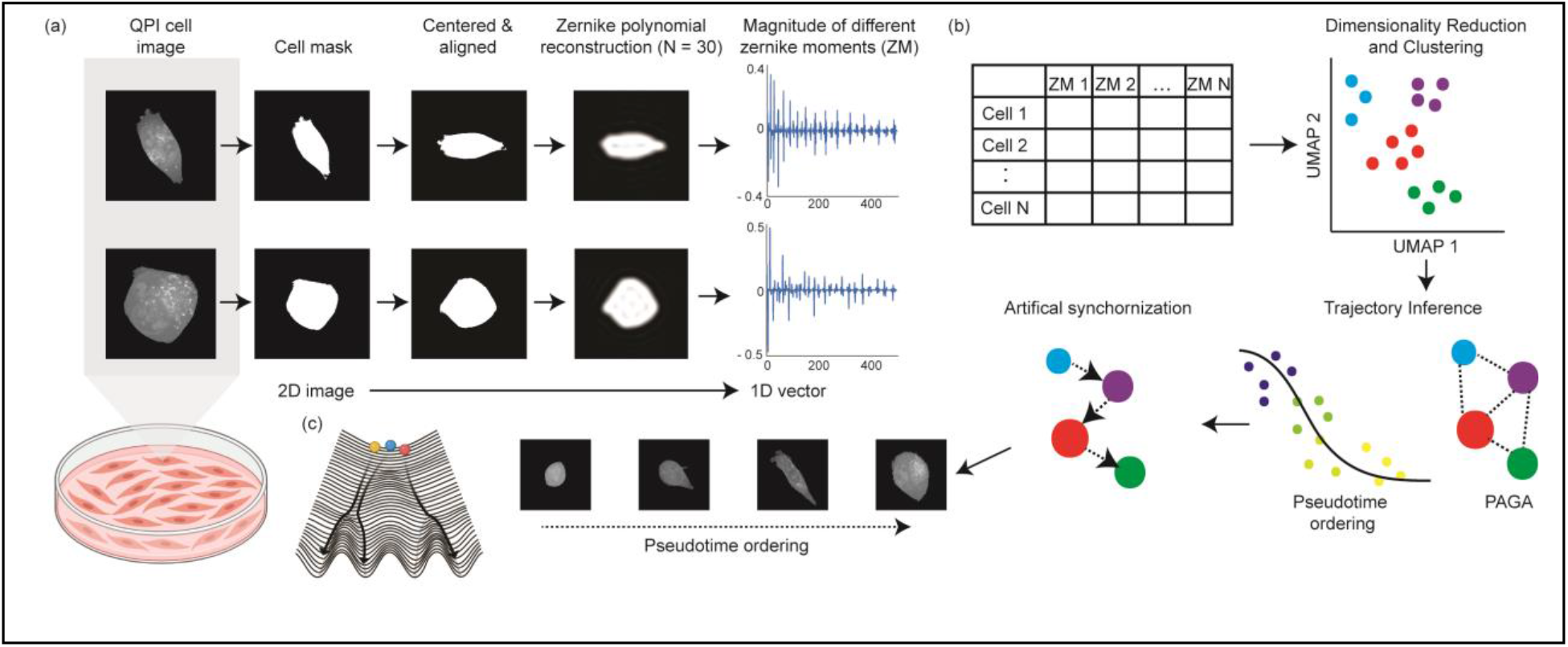
Schematic of the analytical pipeline for Cell-TIMP. (a) QPI cell images are taken from the microscope. The cell images are then segmented, and a cell mask is obtained. The mask images are then centered and aligned along its major axis. Thereafter, Zernike polynomial calculation is done to obtain Zernike moments for each basis function. A reconstructed figure shows that N = 30 is sufficient for most cell morphology application. (b) Once we have converted cell morphology in a one-dimensional vector, we implement TI analytical pipeline on the data. The method is rooted in partition-based graph abstraction algorithm (PAGA) which consists of partitioning a K-nearest neighbors (K-NN) graph into a given number of clusters and linking these clusters to form trajectories. The partitioning of graph was done with leiden clustering which allows user to avoid guessing the number of clusters required and UMAP was used for low-dimensional visualization purpose. (c) A schematic depicting the pathway for Waddington landscape plot.

In recent times, there has been an outburst of analytical tools for trajectory inference (TI) in the context of single-cell transcriptomics that highlight the heterogeneity generated within a cell population [33] [34]. Typically, TI methods start with a vector dataset where each row represents value pertaining to each cell. Then, this matrix is subjected to dimensionality reduction and clustering, followed by tools that connect the trajectory between the cells or clusters in an optimized method. TI techniques operate on a foundation of graph-based principles. Each data point is symbolized as a node, interconnected with neighboring nodes through edges that reflect pairwise similarities derived from distance metrics. Within these topological frameworks, continuous pathways emerge, marked by a commencement point (a root), potential branch points, and one or multiple endpoints. By tracing these extremities, we can ascertain the sequence of intermediary points that manifest along these trajectories. This procedure is termed pseudo-temporal ordering.

In this study, we first started with a synthetic dataset where we tried to see how well computer-generated shapes behave when subjected to dimensionality reduction and clustering algorithm [35]. Following the effective implementation of this methodology on the synthetic dataset, we applied it to the Swedish leaf dataset to assess the performance of unsupervised clustering against supervised labels, confirming the reliability of our analytical pipeline. Additionally, this comparison functions as a vital validation step, which is particularly important when integrating an analytical tool originally devised for disparate objectives. Thereafter, we apply our Zernike preprocessing steps combined with TI tools to classify cell types based on cluster similarity for Leukemia and then order them on a pseudo-time trajectory based on their morphological parameters, which reflects their differentiation pathway. Finally, we applied this analytical pipeline to a breast cancer dataset. In cases where extracting meaningful insights from supervised labels on a low-dimensional plot proved challenging, our approach demonstrates a clear progression in the trajectory of cancer evolution: from parental to circulating to metastatic states based on cellular morphology alone.

## Results and discussions

TI analysis tool works on one-dimensional vector data. Therefore, our first challenge was to convert two-dimensional morphology to a one-dimensional vector. To solve this issue, we used Zernike polynomials on the cellular mask [36] [37]. TI tools can only work properly when we have a well-behaved normalized matrix [33] [34]. If we combine too many dimensions and dimensionless geometrical parameters, the trajectory inference can be erroneous since each matrix column has a different scaling range and cannot be properly row-normalized for each cell. We selected the method of using the Zernike polynomial for its ability to faithfully reconstruct cell shapes while maintaining scale and rotational invariance [38]. Since the resultant Zernike moments are calculated rather than analyzed through a data-driven approach, the resultant data matrix is well-behaved and normalized. This has resulted in utilizing Zernike polynomials in several fields dealing with shape, character, and face recognition [38] [39]. It has been particularly important in the field of optics in defining optical aberration of the eye [40] and optical instrument testing and design [41]. It has also been previously used for single-cell image analysis [42] [43] [44]; however, its utilization has been limited to seeing clusters in low-dimensional plots and classification problems. QPI (Quantitative Phase Imaging) imaging stands as a successor to phase contrast microscopy, an innovation credited to Frits Zernike; therefore, it is almost poetic that in the downstream analysis of these phase images of cells, Zernike polynomials can play such an instrumental role. The workflow for the analysis is shown in Figure 1.

### Analysis of synthetic dataset

We first applied our analytical pipeline to a synthetic dataset which was generated by the implementation of bezier curves [45] [46]. A Bezier curve is a parametric curve where a set of discrete control points defines a smooth curve intending to imitate a shape. In this work, we used these curves to form a closed shape that mimics cell morphology. This was done by having control points 3 and 4 as less than 3 points lead to straight lines, and more than 4 points can cause the shape to overlap and loop, which is unwanted. To form a smooth, closed shape, we made sure they were cyclic, i.e., the transition between the start and end point was smooth. A detailed code implementation can be found in the materials and methods section. Some sample images of the shapes are shown in Figure 2a.

**Figure 2:**
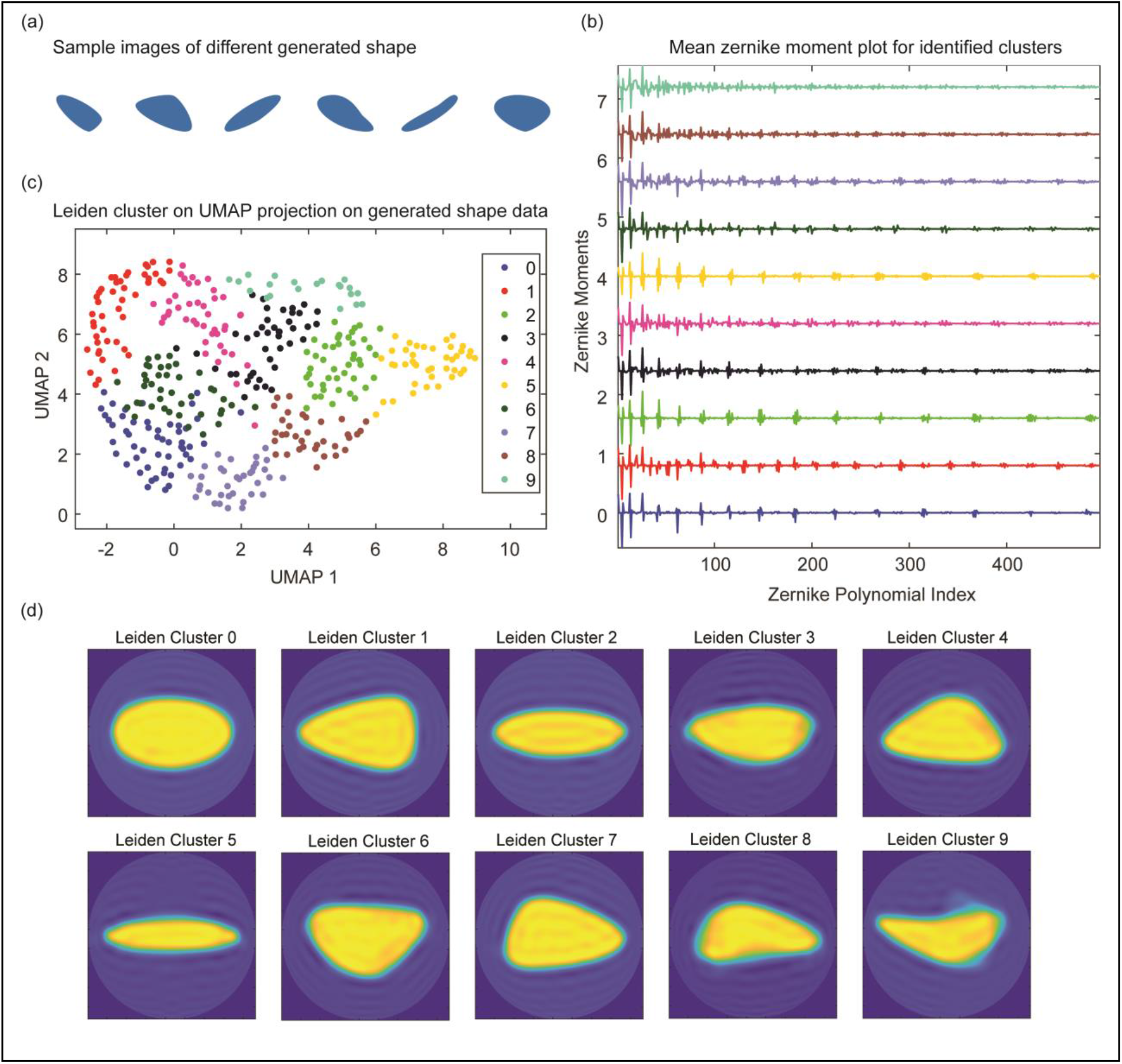
Morphological analysis on computer generated shapes. (a) Sample shapes created by 3 and 4 points bezier curves. (b) Spectra plot of mean Zernike moments for identified clusters for different shapes. (c) Scatter plot on a UMAP representation highlighting different identified clusters through Leiden clustering. (d) Reconstructed shape from average Zernike moment value of identified cluster.

After generating the shape, we generated a mask and centered the image. We also aligned the image along the major axis of the shape as a pre-processing step. The image was then converted into a polar coordinate system (r, ɵ) from a cartesian system. After that, we used Zernfun [47] to compute the Zernike functions Z_nm_ (r, ɵ). The output matrix is a column matrix with values for every (n,m) pair and one row for every (r, ɵ) pair. From here, we get the values of the Zernike moment. The average value of the Zernike moment for each identified cluster is plotted in Figure 2b, and the reconstructed shape from the mean values of each cluster is plotted in Figure 2d.

In Figure 2c, we show different shapes on a low-dimensional plot. In this plot, we see that similar structures are spatially closer in the plot, and structures with less similarity are further apart. The average shape of cluster 0 differs greatly from cluster 5; hence, they are placed far apart along the x-axis. We see a gradual change of shape when we move from cluster 1 to cluster 7, with clusters 4 and 6 along the way. This path follows a gradual and subtle change in shape, encompassing all the dimensions and dimensionless parameters. A similar remark can be made for a path from cluster 9 to 8 with clusters 2 and 3 on the way, highlighting the change in concavity of the shape. Seeing the ability of the analytical method to cluster similar-looking shapes together motivated us to apply this method to real-world supervised datasets.

### Analysis of Swedish leaf dataset

The Swedish leaf dataset [48] is a standardized supervised labeled dataset which has been used earlier in the field of computer vision problems pertaining to shape recognition. The dataset contains images of 15 different classes, with each class having 75 images. For our work, we omitted two classes from the dataset, namely - Sorbus aucuparia and Sorbus intermedia. These two classes were omitted from our study as they had multiple leaves in a single field of view (fern-like structure). We implemented our analytical framework on the remaining 13 classes namely - Ulmus carpinifolia, Acer, Salix aurita, Quercus, Alnus incana, Betula pubescens, Salix alba ‘Sericea’, Populus tremula, Ulmus glabra, Salix sinerea, Populus, Tilia, Fagus sylvatica. The analytical process was similar to the previous case, where we first created the mask, centered and aligned the image, and then computed the Zernike moments for orthogonal polynomials in the polar coordinate. Figure 3a shows a UMAP plot with supervised label where we see that some distinct leaf shapes like Salix Alba, Populus tremula, Acer, Alnus incana, are clustered appropriately while for some labels like Ulmus carpinifolia, Salix aurita, Ulmus glabra, the dots are overlapping. Such heterogeneity and overlap are also common during the cell differentiation process, and therefore, in order to properly define such heterogeneity, we performed unsupervised clustering with Leiden clusters shown in Figure 3b.

**Figure 3:**
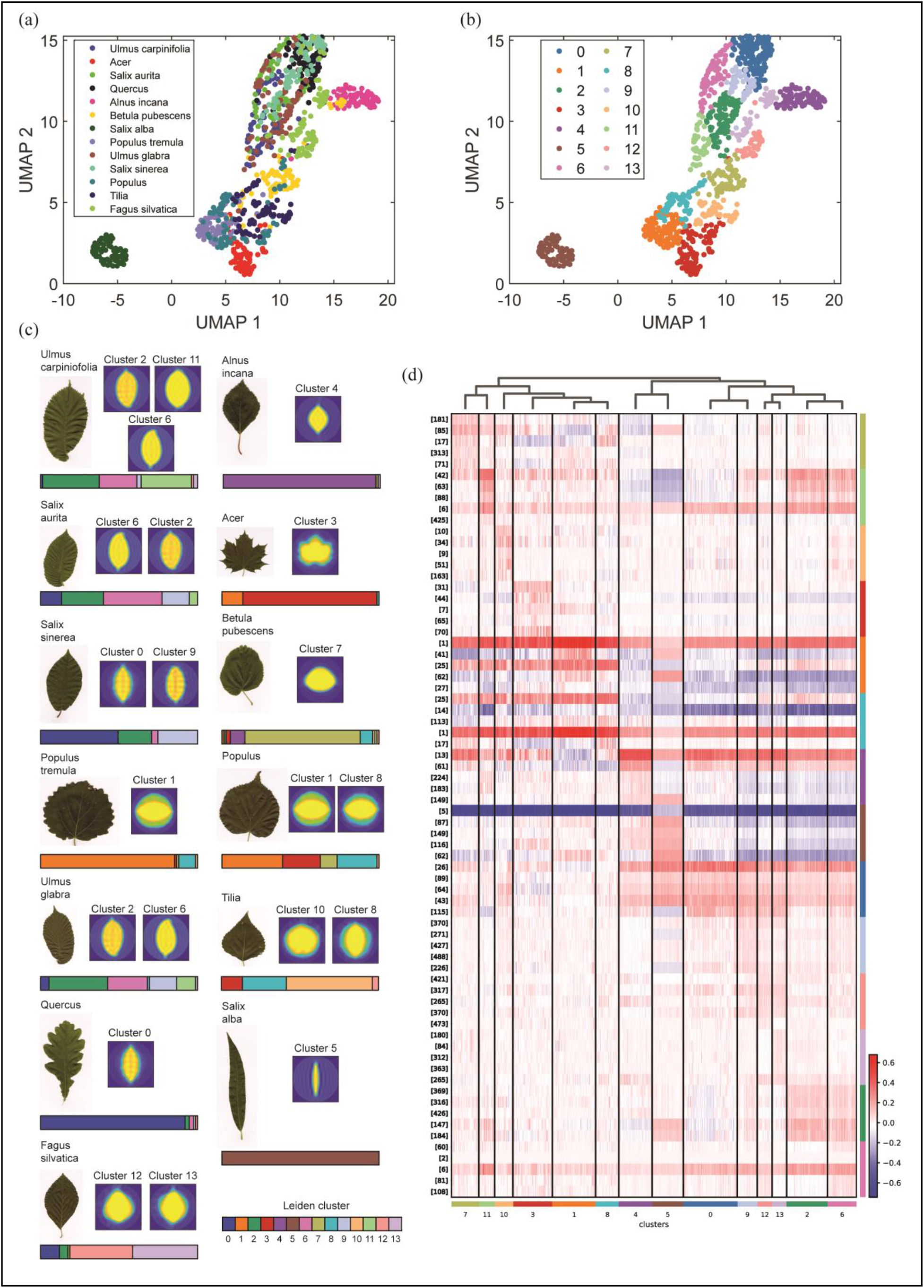
Morphological analysis on Swedish leaf dataset. (a) UMAP plot with actual label after converting different leaf shapes into Zernike polynomials. (b) UMAP plot with unsupervised leiden clusters. (c) Plot showing sample leaf image for each label and the average reconstructed shapes of dominant leiden clusters. Below each figure, we have a colored stacked bar plot for percentage contribution of each leiden cluster constituting the given leaf. (d) Heatmap of top 5 variables from each cluster which were responsible for identification of clusters. The variable number on the left stands for Zernike polynomial index and each row corresponds to the value of Zernike moment for the given polynomial for all the image in the dataset. The solid black line along the y-axis is the margin for each cluster.

In Figure 3c, we have plotted sample leaf images for each label and the average reconstructed shapes of dominant Leiden clusters which make up that class. Below each figure, we have a colored stacked bar plot for the percentage contribution of each Leiden cluster constituting the given leaf. Almost all the contributions come from a single Leiden cluster for distinct shapes like Alnus incana, Acer, Betula pubescens, Populus tremula, Quercus, and Salix alba. The similarity in the average reconstructed shape with the sample leaf image is also noticeable in these cases. This highlights the ability of our analytical pipeline to distinguish different shapes. For leaves such as Populus and Fagus silvatica, we find that it is usually a combination of two identified supervised clusters that best describes their shape. For a more challenging set, which comprises Ulmus carpinifolia, Salix aurita, and Ulmus glabra leaves, we notice that the representative clusters are also similar for these cases. However, their distributions vary. This is very encouraging on two fronts. First, it ensures that our model is not overfitting, which is a characteristic of unsupervised labeling, and second, even when the shape looks similar, the differences can be picked up when doing a population-level analysis. Apart from clustering, the TI analytical toolbox also allows us to arrange them in hierarchical order, establishing cluster similarity. It also helps us identify which variables were responsible for differentiating the clusters from other shapes. This was achieved through scanpy [49] toolbox in python. In Figure 3d, we have a hierarchical arrangement of identified clusters and a heatmap of the top 5 variables from each identified cluster, which were instrumental in differentiating them from other clusters. The variable number on the left stands for the Zernike polynomial index, and each pixel in a particular row corresponds to the value of the Zernike moment corresponding to the polynomial index. The heatmap helps us track the subtle changes in a quantitative manner. It also helps us identify the important variables that can further help in a light-weight shape prediction model.

### Analysis of leukemia dataset

To show the efficacy of the developed analytical pipeline on a cellular dataset, we used leukemic B cells from early (REH, RS4;11) and late (MN60) stages of malignancy along with healthy B-cells (Control) for morphological characterization as shown in figure 4a. The QPI images were acquired by suspending 10-15 uL of cell suspensions from cultures at a density of 10^6^ cells/mL sandwiched between coverslips separated by a secure seal spacer (Invitrogen) [27]. We employed diffraction phase microscopy of the 3M system described previously for acquiring interferograms [50]. The raw interferograms were processed using the inverse Fourier transform method to reveal phase images. Like previous cases, we first created the mask from phase images, then centered and aligned the image, and finally computed the Zernike moments for orthogonal polynomials in the polar coordinate.

**Figure 4:**
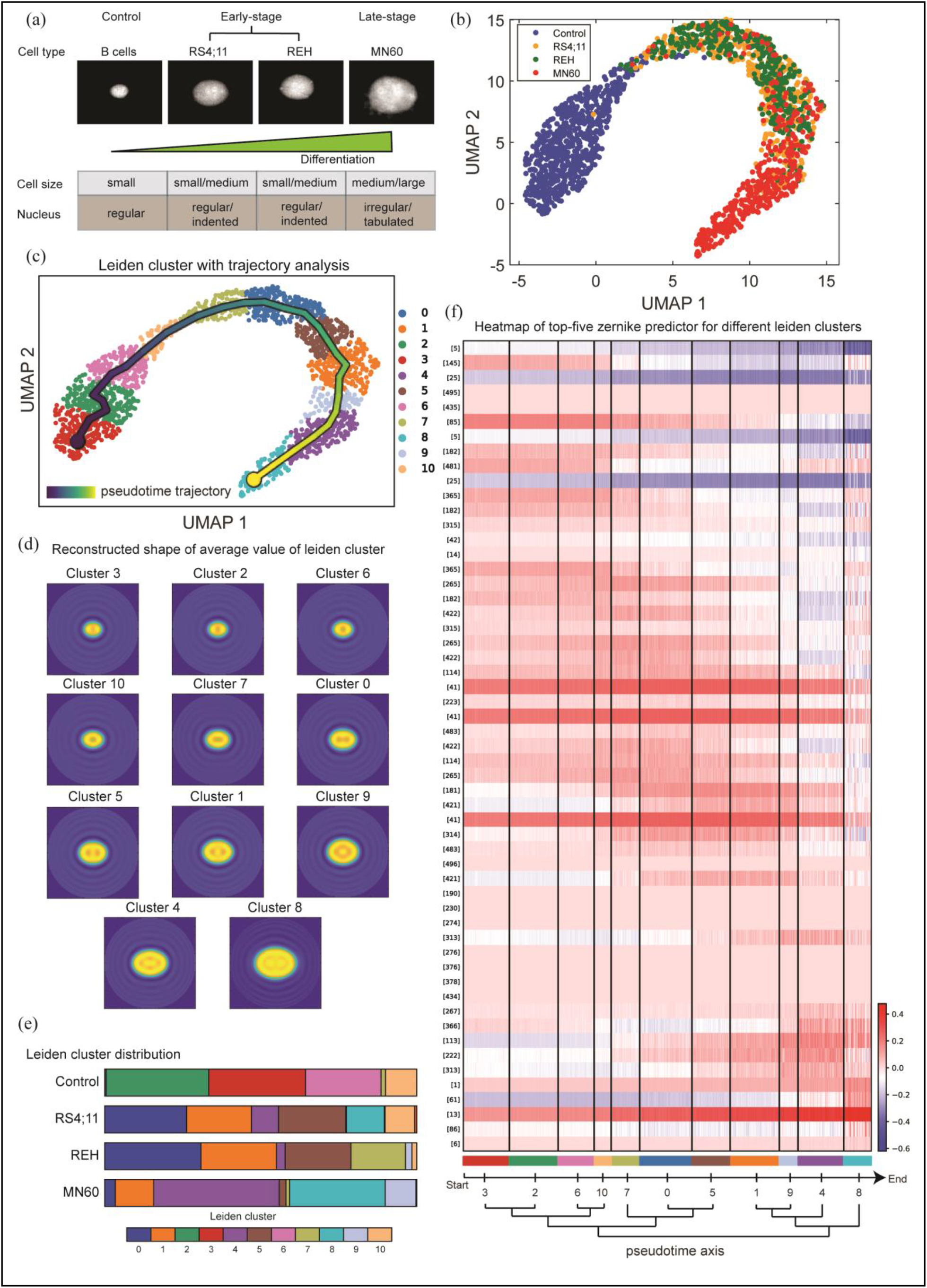
Morphological analysis on leukemia dataset. (a) Sample images of healthy B-cells (Control), early (REH, RS4;11) and late (BALL-1, MN60) stages of malignancy. (b) UMAP plot with actual label after converting cell shape into Zernike polynomials. (c) UMAP plot with unsupervised clustering labels. (d) Reconstructed shape from average value of computed Zernike polynomials from each cluster. (e) Leiden cluster distribution for each supervised label showing the majority of clusters along the differentiation timepoint align well with the pseudotime axis. (f) Heatmap of top 5 variables from each cluster which were responsible for identification of clusters that are aligned on a pseudotime axis. The variable number on the left stands for Zernike polynomial index and each row corresponds to the value of Zernike moment for the given polynomial for all the images in the dataset. The solid black line along the y-axis is the margin for each cluster.

Figure 4b shows the UMAP plot after analyzing the matrix created by Zernike moments. In this plot, we see clear segregation between the control and early stages, and there is some overlap between the early and late stages. When comparing RS4:11 and REH, both of which constitute the early stages, there isn’t clear segregation. This is supported by existing literature where it has been shown that these cell types are very similar in morphology [51] [52]. Next, we show a UMAP plot with unsupervised Leiden clustering in Figure 4c. Knowing that leukemia progression starts from healthy cells, we identified cluster 3 as our starting point. Thereafter, a trajectory was created using scfates python library [53]. The trajectory construction by PAGA (Partition-based graph abstraction) led to the final representation, displaying the ordered sequence of the Leiden clusters in Figure 4c. Comparing the two UMAP plots, we see that several clusters coexist at each differentiation kinetic time, implying shape heterogeneity of the process (Figure 4e). The mean reconstructed shape from each cluster is plotted in Figure 4d, showing the gradual evolution of cell shape from healthy to malignant. To visualize the changes associated with important Zernike polynomials, a heatmap was constructed by plotting the Zernike moment of the top 5 polynomials for each cluster in order of importance against the pseudo-time axis (Figure 4f). The gradual changes in the value of Zernike moment along the pseudo-time axis can be seen for several polynomials like 5, 145, 25, 85, and 42. This highlights the power of the method to track subtle changes in morphology qualitatively. Comparing the pseudo-time axis at the bottom of Figure 4f and label-wise cluster distribution in Figure 4e, we see that most clusters constituting the Control cell type are at the beginning of the time axis. The early stages (RS4;11 and REH) have major contributions from clusters in the middle of the time axis, and the late stage (MN60) have major contributions from clusters at the end of the time axis.

### Analysis of breast cancer dataset

To further validate the efficacy of our method, we ran our analysis on breast cancer cell lines with limited cell images (N = 157) (Figure 5a). The cell lines selected represent breast cancer cell lines throughout their lifetime, starting from the primary organ, then as circulating tumor cells, and finally homing to a distant organ, metastatization to the lungs [12]. The three cell lines were seeded in glass coverslip-bottom Petri dishes for tomography images. The imaging was performed on an ODT system (HT-1H, Tomocube Inc., Republic of Korea) comprised of a 60X water-immersion objective (1.2 NA), an off-axis Mach-Zehnder interferometer with a 532 nm laser and a digital micromirror device (DMD) for tomographic scanning of each cell [54].

**Figure 5:**
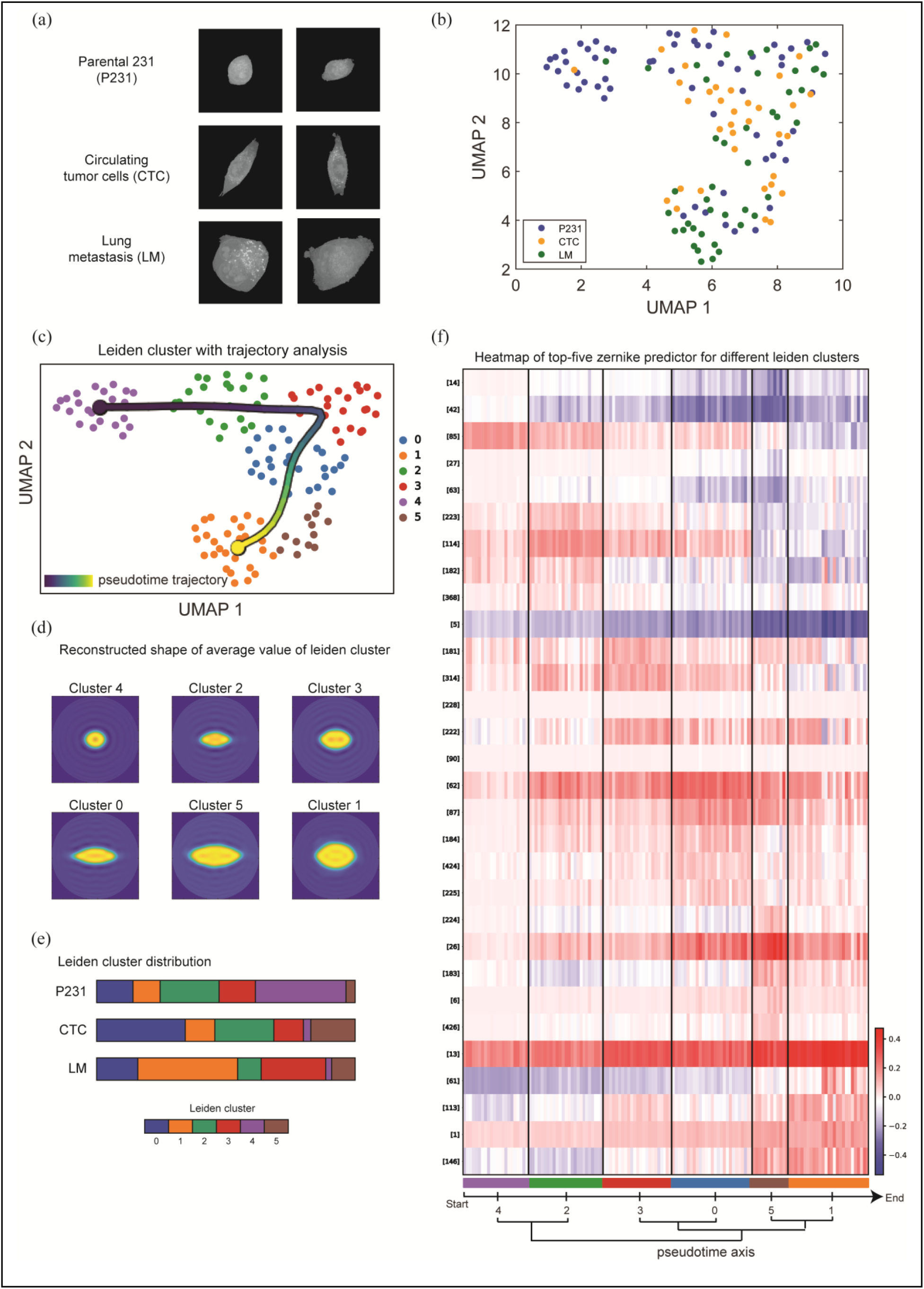
Morphological analysis on breast cancer dataset. (a) Sample QPI images of Parental 231 (P231), Circulating tumor cells (CTC) and Lung Metastasis (LM) cells. (b) UMAP plot with actual label after converting cell shape into Zernike polynomials. (c) UMAP plot with unsupervised clustering labels. (d) Reconstructed shape from average value of computed Zernike polynomials from each cluster. (e) Leiden cluster distribution for each supervised label showing most clusters along the differentiation timepoint align well with the pseudotime axis. (f) Heatmap of top 5 variables from each cluster which were responsible for identification of clusters that are aligned on a pseudotime axis. The variable number on the left stands for Zernike polynomial index and each row corresponds to the value of Zernike moment for the given polynomial for all the images in the dataset. The solid black line along the y-axis is the margin for each cluster.

The shape analysis methods that are based on neural networks or other data-driven methods are usually not able to give meaningful results with such a small number of images [55]. In our previous work [12], we have reported that while quantitative phase images show subtle but significant morphological differences, they are insufficient for robust classification of closely related cells at a single-cell analytical resolution, particularly when the training data is relatively scarce. The random forest classifier was found to asymptotically plateau around 46% (compared to 33.3% random chance).

This is also evident in Figure 5b, where the UMAP plot with supervised labels is not able to provide any significant insight into these cell lines. This is directly a result of continual and asynchronous movement between cellular states leading to heterogeneity, and in smaller datasets, conventional analytical methods leave us blind without revealing much. Our analysis methods first involved unsupervised clustering using the Leiden cluster and then PAGA-based trajectory construction, as described above in Figure 1. By plotting the reconstructed shape of the average value of the Leiden cluster, we were able to attach physical meaning to the constructed trajectory (Figure 5c and 5d). The cells start as small, rounded structures and gradually get bigger and slender. The slender shape is known to enhance migration potential [4] [56] [57]. And finally, they get bigger in shape while losing their slenderness [58]. When we look at the supervised label with the percent distribution of identified Leiden clusters that constitute them (Figure 5e), we see that P231 cells, which are primary cancer cells, have most of the contribution from clusters 4 and 2, which are early on the pseudo-time axis (Figure 5f). These cell shapes are mostly oval, with some cells being more directional and circular than others. Thereafter, the majority of contribution for CTC cells comes from clusters 0 and 2, which are in the middle of the pseudo-time axis. During metastasis, the migrating cancer cell has to make its way through the extracellular matrix and then flow into the bloodstream; therefore these microenvironment constraints promote the cell shape to become more slender and spindle-shaped [56] [57]. Finally, most of the contribution for LM is coming from cluster 1, which is at the end of the time axis. These cells have metastasized to a distant organ and are shedding away its migration potential-related properties. However, they are now much bigger compared to other cells as they are more mature [58]. Interestingly, some of the contributions for LM come from cluster 3 as well, which, although in the middle, reflects a cell shape that is transitioning from the rounder morphology to spindle-like morphology. Figure 5f also shows a heatmap of the top 5 polynomials for each cluster in order of importance against the pseudo-time axis, providing quantitative details of morphological differences between different clusters.

## Conclusion

In summary, our work demonstrates that the use of TI tools, which have been initially introduced for sc-RNA-seq analysis, could be beneficial to be employed for morphological analysis. Since there is no manual input required at any step of the analysis, this methodology is completely automated, and when combined with high-throughput phase imaging, it paves the way for a novel generation of morphological phenotyping by clearly accounting for cellular heterogeneity.

## Materials and Methods

### Synthetic dataset

Synthetic cell shape was created using beizer curves implementation in python. This implementation can be found at: https://stackoverflow.com/questions/50731785/create-random-shape-contour-using-matplotlib

### Leaf dataset

The leaf dataset is taken from [48]. This dataset can be found at: https://www.cvl.isy.liu.se/en/research/datasets/swedish-leaf/

### Cell culture protocol for Leukemia cells

We purchased the early-stage leukemia cell lines REH and RS4;11 from American Type Culture Collection (ATCC, USA) and procured the late-stage leukemia cell lines MN60 from DSMZ (Germany). RS4;11 and REH cells were grown in medium consisting of RPMI-1640, 10% (v/v) fetal bovine serum (FBS), 50 mg/mL penicillin-streptomycin, and 2 mM l-glutamine (Thermo Fisher, Waltham, Massachusetts). The MN60 cell line was cultured in minimum essential medium supplemented with 10% (v/v) FBS, 50 mg/mL penicillin–streptomycin, and 2 mM l-glutamine. All cells were cultured at a density of 10^6^ cells/mL prior to taking them out for measurements.

For healthy control samples, fresh blood samples from three anonymous healthy adult donors were purchased from StemCell Technologies (Vancouver, Canada) and all the experiments were conducted within 24 hours of blood donation. The purchased blood samples contained ethylenediaminetetraacetic acid (EDTA) as an anticoagulant. The B cells were isolated from fresh blood samples using negative selection kits from Stemcell Technologies (EasySep Direct Human B Cell Isolation Kit).

### Cell culture protocol for Breast Cancer cells

An isogenic panel of varying metastatic potential derived from the human breast cancer cell line MDA-MB-231 was used in this study. In addition to the parental MDA-MB-231 cells (P231), the panel consisted of CTC and LM cells previously obtained after orthotopic implantation of the parental cells in the fourth right mammary fat pad of female athymic nu/nu female mouse (NCI) as detailed in our previous publications [59] [60]. The three cell lines were cultured in RPMI-1640 media supplemented with 10% fetal bovine serum (FBS), 100 U/ml penicillin, and 100 μg/ml streptomycin and maintained at 37 °C and 5% CO_2_ in a humidified incubator.

## Acknowledgment

We would like to thank Dr. Rishikesh Pandey for providing us with Leukemia cell images and Dr. Kristine Glunde for providing the breast cancer cell lines.

We acknowledge support from the Air Force Office of Scientific Research (FA9550–22-1–0334) and National Institute of General Medical Sciences (1R35GM149272).

Figure 1 was partially created with Biorender.com.

## References

[1] P. Rangamani et al., “Decoding Information in Cell Shape,” Cell, vol. 154, no. 6, pp. 1356–1369, Sep. 2013, doi: 10.1016/j.cell.2013.08.026.

[2] H.-G. Döbereiner, B. Dubin-Thaler, G. Giannone, H. S. Xenias, and M. P. Sheetz, “Dynamic Phase Transitions in Cell Spreading,” Phys. Rev. Lett., vol. 93, no. 10, p. 108105, Sep. 2004, doi: 10.1103/PhysRevLett.93.108105.

[3] Y. Xiong et al., “Mechanisms Controlling Cell Size and Shape during Isotropic Cell Spreading,” Biophys. J., vol. 98, no. 10, pp. 2136–2146, May 2010, doi: 10.1016/j.bpj.2010.01.059.

[4] A. Mogilner and K. Keren, “The Shape of Motile Cells,” Curr. Biol., vol. 19, no. 17, pp. R762–R771, Sep. 2009, doi: 10.1016/j.cub.2009.06.053.

[5] P.-H. Wu et al., “Evolution of cellular morpho-phenotypes in cancer metastasis,” Sci. Rep., vol. 5, no. 1, p. 18437, Dec. 2015, doi: 10.1038/srep18437.

[6] W.-C. Chen et al., “Functional interplay between the cell cycle and cell phenotypes,” Integr. Biol., vol. 5, no. 3, pp. 523–534, Mar. 2013, doi: 10.1039/c2ib20246h.

[7] C. Bakal, J. Aach, G. Church, and N. Perrimon, “Quantitative Morphological Signatures Define Local Signaling Networks Regulating Cell Morphology,” Science, vol. 316, no. 5832, pp. 1753–1756, Jun. 2007, doi: 10.1126/science.1140324.

[8] P.-H. Wu et al., “Single-cell morphology encodes metastatic potential,” Sci. Adv., vol. 6, no. 4, p. eaaw6938, Jan. 2020, doi: 10.1126/sciadv.aaw6938.

[9] T. J. Fuchs and J. M. Buhmann, “Computational pathology: Challenges and promises for tissue analysis,” Comput. Med. Imaging Graph., vol. 35, no. 7–8, pp. 515–530, Oct. 2011, doi: 10.1016/j.compmedimag.2011.02.006.

[10] A. H. Beck et al., “Systematic Analysis of Breast Cancer Morphology Uncovers Stromal Features Associated with Survival,” Sci. Transl. Med., vol. 3, no. 108, Nov. 2011, doi: 10.1126/scitranslmed.3002564.

[11] C. McQuin et al., “CellProfiler 3.0: Next-generation image processing for biology,” PLOS Biol., vol. 16, no. 7, p. e2005970, Jul. 2018, doi: 10.1371/journal.pbio.2005970.

[12] S. K. Paidi, V. Shah, P. Raj, K. Glunde, R. Pandey, and I. Barman, “Coarse Raman and optical diffraction tomographic imaging enable label-free phenotyping of isogenic breast cancer cells of varying metastatic potential,” Biosens. Bioelectron., vol. 175, p. 112863, Mar. 2021, doi: 10.1016/j.bios.2020.112863.

[13] N. Petrović, G. Moyà-Alcover, A. Jaume-i-Capó, and M. González-Hidalgo, “Sickle-cell disease diagnosis support selecting the most appropriate machine learning method: Towards a general and interpretable approach for cell morphology analysis from microscopy images,” Comput. Biol. Med., vol. 126, p. 104027, Nov. 2020, doi: 10.1016/j.compbiomed.2020.104027.

[14] U. S. Schwarz and S. A. Safran, “Physics of adherent cells,” Rev. Mod. Phys., vol. 85, no. 3, pp. 1327– 1381, Aug. 2013, doi: 10.1103/RevModPhys.85.1327.

[15] J. Ge et al., “Standard fluorescent imaging of live cells is highly genotoxic,” Cytometry A, vol. 83A, no. 6, pp. 552–560, Jun. 2013, doi: 10.1002/cyto.a.22291.

[16] E. C. Jensen, “Use of Fluorescent Probes: Their Effect on Cell Biology and Limitations,” Anat. Rec. Adv. Integr. Anat. Evol. Biol., vol. 295, no. 12, pp. 2031–2036, Dec. 2012, doi: 10.1002/ar.22602.

[17] Y. Park, C. Depeursinge, and G. Popescu, “Quantitative phase imaging in biomedicine,” Nat. Photonics, vol. 12, no. 10, pp. 578–589, Oct. 2018, doi: 10.1038/s41566-018-0253-x.

[18] K. Kim, K. S. Kim, H. Park, J. C. Ye, and Y. Park, “Real-time visualization of 3-D dynamic microscopic objects using optical diffraction tomography,” Opt. Express, vol. 21, no. 26, p. 32269, Dec. 2013, doi: 10.1364/OE.21.032269.

[19] G. Popescu, Y. Park, W. Choi, R. R. Dasari, M. S. Feld, and K. Badizadegan, “Imaging red blood cell dynamics by quantitative phase microscopy,” Blood Cells. Mol. Dis., vol. 41, no. 1, pp. 10–16, Jul. 2008, doi: 10.1016/j.bcmd.2008.01.010.

[20] S. Tanwar et al., “Targeted Enzyme Activity Imaging with Quantitative Phase Microscopy,” Nano Lett., vol. 23, no. 10, pp. 4602–4608, May 2023, doi: 10.1021/acs.nanolett.3c01090.

[21] P. Anantha, Z. Liu, P. Raj, and I. Barman, “Optical diffraction tomography and Raman spectroscopy reveal distinct cellular phenotypes during white and brown adipocyte differentiation,” Biosens. Bioelectron., vol. 235, p. 115388, Sep. 2023, doi: 10.1016/j.bios.2023.115388.

[22] Y. R. He et al., “Cell Cycle Stage Classification Using Phase Imaging with Computational Specificity,” ACS Photonics, vol. 9, no. 4, pp. 1264–1273, Apr. 2022, doi: 10.1021/acsphotonics.1c01779.

[23] V. Ayyappan et al., “Identification and Staging of B-Cell Acute Lymphoblastic Leukemia Using Quantitative Phase Imaging and Machine Learning,” ACS Sens., vol. 5, no. 10, pp. 3281–3289, Oct. 2020, doi: 10.1021/acssensors.0c01811.

[24] N. Goswami et al., “Label-free SARS-CoV-2 detection and classification using phase imaging with computational specificity,” Light Sci. Appl., vol. 10, no. 1, p. 176, Sep. 2021, doi: 10.1038/s41377-021-00620-8.

[25] G. Wali, S. Berkovsky, D. R. Whiten, A. Mackay-Sim, and C. M. Sue, “Single cell morphology distinguishes genotype and drug effect in Hereditary Spastic Paraplegia,” Sci. Rep., vol. 11, no. 1, p. 16635, Aug. 2021, doi: 10.1038/s41598-021-95995-4.

[26] A. Marconi et al., “In Vivo Melanoma Cell Morphology Reflects Molecular Signature and Tumor Aggressiveness,” J. Invest. Dermatol., vol. 142, no. 8, pp. 2205–2216.e6, Aug. 2022, doi: 10.1016/j.jid.2021.12.024.

[27] S. K. Paidi et al., “Raman and quantitative phase imaging allow morpho-molecular recognition of malignancy and stages of B-cell acute lymphoblastic leukemia,” Biosens. Bioelectron., vol. 190, p. 113403, Oct. 2021, doi: 10.1016/j.bios.2021.113403.

[28] S. J. Altschuler and L. F. Wu, “Cellular Heterogeneity: Do Differences Make a Difference?,” Cell, vol. 141, no. 4, pp. 559–563, May 2010, doi: 10.1016/j.cell.2010.04.033.

[29] M. Tellez-Gabriel, B. Ory, F. Lamoureux, M.-F. Heymann, and D. Heymann, “Tumour Heterogeneity: The Key Advantages of Single-Cell Analysis,” Int. J. Mol. Sci., vol. 17, no. 12, p. 2142, Dec. 2016, doi: 10.3390/ijms17122142.

[30] Y. Tian et al., “Single-cell transcriptomic profiling reveals the tumor heterogeneity of small-cell lung cancer,” Signal Transduct. Target. Ther., vol. 7, no. 1, p. 346, Oct. 2022, doi: 10.1038/s41392-022-01150-4.

[31] C. H. Waddington, The Strategy of the Genes, 0 ed. Routledge, 2014. doi: 10.4324/9781315765471.

[32] J. M. Phillip, K.-S. Han, W.-C. Chen, D. Wirtz, and P.-H. Wu, “A robust unsupervised machine-learning method to quantify the morphological heterogeneity of cells and nuclei,” Nat. Protoc., vol. 16, no. 2, pp. 754–774, Feb. 2021, doi: 10.1038/s41596-020-00432-x.

[33] M. D. Luecken and F. J. Theis, “Current best practices in single-cell RNA-seq analysis: a tutorial,” Mol. Syst. Biol., vol. 15, no. 6, p. e8746, Jun. 2019, doi: 10.15252/msb.20188746.

[34] W. Saelens, R. Cannoodt, H. Todorov, and Y. Saeys, “A comparison of single-cell trajectory inference methods,” Nat. Biotechnol., vol. 37, no. 5, pp. 547–554, May 2019, doi: 10.1038/s41587-019-0071-9.

[35] V. A. Traag, L. Waltman, and N. J. Van Eck, “From Louvain to Leiden: guaranteeing well-connected communities,” Sci. Rep., vol. 9, no. 1, p. 5233, Mar. 2019, doi: 10.1038/s41598-019-41695-z.

[36] V. F. Zernike, “Beugungstheorie des schneidenver-fahrens und seiner verbesserten form, der phasenkontrastmethode,” Physica, vol. 1, no. 7–12, pp. 689–704, May 1934, doi: 10.1016/S0031-8914(34)80259-5.

[37] K. Niu and C. Tian, “Zernike polynomials and their applications,” J. Opt., vol. 24, no. 12, p. 123001, Dec. 2022, doi: 10.1088/2040-8986/ac9e08.

[38] A. Khotanzad and Y. H. Hong, “Invariant image recognition by Zernike moments,” IEEE Trans. Pattern Anal. Mach. Intell., vol. 12, no. 5, pp. 489–497, May 1990, doi: 10.1109/34.55109.

[39] C. Singh, E. Walia, and N. Mittal, “Rotation invariant complex Zernike moments features and their applications to human face and character recognition,” IET Comput. Vis., vol. 5, no. 5, p. 255, 2011, doi: 10.1049/iet-cvi.2010.0020.

[40] L. N. Thibos, R. A. Applegate, J. T. Schwiegerling, and R. Webb, “Standards for Reporting the Optical Aberrations of Eyes,” J. Refract. Surg., vol. 18, no. 5, Sep. 2002, doi: 10.3928/1081-597X-20020901-30.

[41] V. N. Mahajan, “Zernike Polynomial and Wavefront Fitting,” in Optical Shop Testing, 1st ed., D. Malacara, Ed., Wiley, 2007, pp. 498–546. doi: 10.1002/9780470135976.ch13.

[42] E. Alizadeh, S. M. Lyons, J. M. Castle, and A. Prasad, “Measuring systematic changes in invasive cancer cell shape using Zernike moments,” Integr. Biol., vol. 8, no. 11, pp. 1183–1193, 2016, doi: 10.1039/C6IB00100A.

[43] A. Tahmasbi, F. Saki, and S. B. Shokouhi, “Classification of benign and malignant masses based on Zernike moments,” Comput. Biol. Med., vol. 41, no. 8, pp. 726–735, Aug. 2011, doi: 10.1016/j.compbiomed.2011.06.009.

[44] G. Apostolopoulos, S. Tsinopoulos, and E. Dermatas, “Recognition and identification of red blood cell size using Zernike moments and multicolor scattering images,” in 2011 10th International Workshop on Biomedical Engineering, Kos, Greece: IEEE, Oct. 2011, pp. 1–4. doi: 10.1109/IWBE.2011.6079035.

[45] G. Farin, Curves and Surfaces for CAGD: A Practical Guide. Morgan-Kaufmann, 2022.

[46] Pomax, A Primer on Bézier Curves. [Online]. Available: https://pomax.github.io/bezierinfo/

[47] Fricker, Paul, “Zernike Polynomials.” MATLAB Central File Exchange, 2023. [Online]. Available: https://www.mathworks.com/matlabcentral/fileexchange/7687-zernike-polynomials

[48] O. Söderkvist, “Computer Vision Classification of Leaves from Swedish Trees,” Linkopping University, 2001. [Online]. Available: https://liu.diva-portal.org/smash/record.jsf?pid=diva2%3A303038&dswid=9519

[49] F. A. Wolf, P. Angerer, and F. J. Theis, “SCANPY: large-scale single-cell gene expression data analysis,” Genome Biol., vol. 19, no. 1, p. 15, Dec. 2018, doi: 10.1186/s13059-017-1382-0.

[50] R. Pandey et al., “Integration of diffraction phase microscopy and Raman imaging for label-free morpho-molecular assessment of live cells,” J. Biophotonics, vol. 12, no. 4, p. e201800291, Apr. 2019, doi: 10.1002/jbio.201800291.

[51] H. G. Drexler and R. A. Macleod, “History of leukemia-lymphoma cell lines: History of leukemia-lymphoma cell lines,” Hum. Cell, vol. 23, no. 3, pp. 75–82, Aug. 2010, doi: 10.1111/j.1749-0774.2010.00087.x.

[52] S. Managò et al., “A reliable Raman-spectroscopy-based approach for diagnosis, classification and follow-up of B-cell acute lymphoblastic leukemia,” Sci. Rep., vol. 6, no. 1, p. 24821, Apr. 2016, doi: 10.1038/srep24821.

[53] L. Faure, R. Soldatov, P. V. Kharchenko, and I. Adameyko, “scFates: a scalable python package for advanced pseudotime and bifurcation analysis from single-cell data,” Bioinformatics, vol. 39, no. 1, p. btac746, Jan. 2023, doi: 10.1093/bioinformatics/btac746.

[54] S. Shin, K. Kim, J. Yoon, and Y. Park, “Active illumination using a digital micromirror device for quantitative phase imaging,” Opt. Lett., vol. 40, no. 22, p. 5407, Nov. 2015, doi: 10.1364/OL.40.005407.

[55] C. Bishop, Pattern Recognition and Machine Learning. in Information Science and Statistics. Springer New York, NY.

[56] D. L. Bodor, W. Pönisch, R. G. Endres, and E. K. Paluch, “Of Cell Shapes and Motion: The Physical Basis of Animal Cell Migration,” Dev. Cell, vol. 52, no. 5, pp. 550–562, Mar. 2020, doi: 10.1016/j.devcel.2020.02.013.

[57] K. A. Rejniak, “Circulating Tumor Cells: When a Solid Tumor Meets a Fluid Microenvironment,” in Systems Biology of Tumor Microenvironment, vol. 936, K. A. Rejniak, Ed., in Advances in Experimental Medicine and Biology, vol. 936., Cham: Springer International Publishing, 2016, pp. 93–106. doi: 10.1007/978-3-319-42023-3_5.

[58] S. M. Lyons et al., “Changes in cell shape are correlated with metastatic potential in murine and human osteosarcomas,” Biol. Open, vol. 5, no. 3, pp. 289–299, Mar. 2016, doi: 10.1242/bio.013409.

[59] A. Rizwan et al., “Metastatic breast cancer cells in lymph nodes increase nodal collagen density,” Sci. Rep., vol. 5, no. 1, p. 10002, May 2015, doi: 10.1038/srep10002.

[60] A. Rizwan, S. K. Paidi, C. Zheng, M. Cheng, I. Barman, and K. Glunde, “Mapping the genetic basis of breast microcalcifications and their role in metastasis,” Sci. Rep., vol. 8, no. 1, p. 11067, Jul. 2018, doi: 10.1038/s41598-018-29330-9.

